# Influence of upper limb training and analyzed muscles on estimate of physical activity during cereal grinding using saddle quern and rotary quern

**DOI:** 10.1101/2020.11.26.399600

**Authors:** Michal Struška, Martin Hora, Thomas R. Rocek, Vladimír Sládek

## Abstract

Experimental grinding has been used to study the relationship between human humeral robusticity and cereal grinding in the early Holocene. However, such replication studies raise two questions regarding the robusticity of the results: whether female nonathletes used in previous research are sufficiently comparable to early agricultural females, and whether previous analysis of muscle activation of only four upper limb muscles is sufficient to capture the stress of cereal grinding on upper limb bones. We test the influence of both of these factors. Electromyographic activity of eight upper limb muscles was recorded during cereal grinding in an athletic sample of 10 female rowers and a nonathletic sample of 25 females and analyzed using both an eight- and four-muscle model. Athletes had lower activation than nonathletes in the majority of measured muscles, but most of these differences were non-significant. Furthermore, both athletes and nonathletes had lower muscle activation during saddle quern grinding than rotary quern grinding suggesting that the nonathletic sample can be used to model early agricultural females during saddle and rotary quern grinding.

Similarly, in both eight- and four-muscle models, upper limb loading was lower during saddle quern grinding than during rotary quern grinding, suggesting that the upper limb muscles may be reduced to the previously used four-muscle model for evaluation of the upper limb loading during cereal grinding. Another implication of our measurements is to question the assumption that skeletal indicators of high involvement of the biceps brachii muscle can be interpreted as specifically indicative of saddle quern grinding.

## Introduction

The experimental reconstruction of habitual tasks of past populations is often used to test hypotheses linking skeletal markers of activity with the behavior of past humans [1–4]. Aside from animal models, it is impractical to observe developmental changes in human skeletal morphology as a result of activity patterns; thus, measurements of muscle activity in modern subjects replicating past activities serve as a route for inferring the impact of prehistoric behaviors. Two significant issues underlie such experimental replicative experiments: the choice of appropriate modern experimental subjects, and the choice of which muscle activity to measure.

The first issue raises the question of the applicability of experiments using modern sedentary and physically inactive populations whose lifelong pattern of physical exertion differs drastically from the heavy physical workload of the prehistoric populations of interest. These modern groups are typically assumed to be inappropriate for such experimental comparison with prehistoric samples, and instead studies of modern athletes are often considered a starting point for such research (e.g. [5–7]). However, while modern athletes probably do more closely resemble the physical condition of most prehistoric groups, here we consider whether comparisons among the physical demands of various prehistoric physical activities may not require restriction of the modern test sample to trained athletes if the *relative* differences between the activities are reflected in *relative* differences between experimental groups, even non-athletic ones.

A similar question relates to the choice of muscular data needed to conduct comparative studies of alternate activity patterns. Ideally, assessments of musculoskeletal consequences of physical exertion should measure as many as possible of the muscles that are potentially recruited during the task. However, experimental reconstruction using electromyography (EMG) often analyzes a limited set of muscles [3,4]. The number of muscles analyzed in experimental reconstructions might be limited by the number of available sensors and the duration of experimental sessions. Reducing the set of analyzed muscles might be inevitable, but the effect of reducing the investigated upper limb muscles to a small set has not been explored. Thus, it is useful to examine if broad sampling of muscle activity is necessary, or whether smaller selective samples of relevant major muscles can provide equivalent information. Finally, when consistent results are found, we can compare these against skeletal inferences of past activity patterns.

In this study we examine these two questions in the context of research on muscle activation associated with alternative patterns of cereal grinding, comparing reciprocal grinding using a saddle quern to rotary grinding with a rotary quern. Cereal grinding has previously been experimentally reconstructed to estimate the effectiveness of early agricultural grinding tools [4], the influence of different grains on the resulting product [8], and upper limb loading during grinding [4]. The pattern of upper limb loading during grinding was used to explain changes in humeral robusticity asymmetry among humans in the early Holocene [4,9]. This research was based on experimental grinding activities by a sample of modern, nonathletic volunteers.

Two recent papers on experimental approaches to prehistoric activity patterns have strengthened the argument for the use of modern athletes specialized in sports involving motions similar to the reconstructed archaeological tasks [5,6]. The influence of a modern athletic sample on experimental results was shown in [6], where athletes were able to throw replicas of Middle Paleolithic spears farther than previously believed based on a wider range of subjects. On the other hand, we don’t know what influence an athletic sample would have on the previous experimental reconstruction of other tasks, such as spear thrusting [3] or cereal grinding [4]. Even less clear is the influence of different levels of athletic training on the *relative* patterns of muscle use or effectiveness during these alternative grinding activities.

Previous analysis of muscle activation during cereal grinding used nonathletic female subjects [4]. However, early Holocene females had extensive experience with grinding, which today’s females lack. Experimental and ethnographic evidence suggest that early Holocene females spent 1.5–7 h/day grinding on saddle quern or 1.3 h/day grinding on rotary quern [4,8,10,11]. Daily repetition of grinding movement in early Holocene females could have had effects on muscle activation, as observed in contemporary manual workers [12–14]. Early agricultural females involved in daily repetition of cereal grinding could have activated their muscles less or could have activated different muscles than modern females that are less involved in upper limb tasks. Therefore, skeletal adjustments in early agricultural females could occur in different places or to lower extent than inferred from nonathletic contemporary females. To account for experience, grinding could be studied in populations that still use manual grinding tools [10,11,15–18], but in those populations, saddle and rotary querns are not used concurrently. Therefore, females from contemporary populations would not have experience with grinding on both saddle and rotary querns. As an alternative, the subsistence behavior of past populations could be studied in athletes, as was recently reviewed for reconstructions of other past human activities [5].

Cereal grinding does not have direct equivalent in any athletic discipline, but there are characteristics in which habitual grinding of early Holocene females resembles rowing. In both grinding and rowing, the upper limbs are involved in repetitive movement pattern. Duration of a rowing cycle can be as low as 1.6 s [19], which is similar to the 1.4 s cycle used for rotary quern grinding in a previous experimental study [4]. Professional rowers spend 11.2 h/week rowing [20], which is close to the estimated time for rotary quern grinding (9.1 h/week) [4]. These similarities between rowers and grinding females suggest that rowers might help us understand the effect of cereal grinding on human muscle activity better than the previously used nonathletic females.

In addition to the question of the appropriate modern population sample for experimental testing is the issue of the choice of muscles to measure. The relationship between cereal grinding and muscle activation during grinding was previously measured in four muscles with insertion on the upper limb (anterior deltoid, infraspinatus, pectoralis major, and long head of the triceps brachii) [4]. While the previous study selected four prime movers of the upper limbs, more muscles may be necessary to see the effect of muscle forces that control shoulder and elbow motions during cereal grinding. The muscles examined in the previous study are mainly responsible for shoulder flexion and rotation and elbow extension [4]. By adding the muscles with other functions, such as shoulder extension and elbow flexion, we may estimate upper limb loading during a greater portion of shoulder movements.

Furthermore, the analysis of the activation of more upper limb muscles would indicate in which muscle insertions we may expect entheseal changes associated with cereal grinding on saddle and rotary quern. This could help connect entheseal changes with changes of grinding technology during adoption and intensification of agriculture. In addition, we could test the activation of the muscles that were suggested to be active during cereal grinding based on entheseal changes, such as the deltoid or biceps brachii during saddle quern grinding [21]. Thus, measures of muscle activation may assist in osteological interpretations of prehistoric activities.

As the first goal, we aim to compare muscle activation during grinding in athletes and nonathletes using electromyography. We expect athletes to have lower muscle activation during grinding than nonathletes. The second goal is to compare upper limb loading during saddle and rotary quern grinding estimated using the eight- and four-muscle models. We expect the eight-muscle model to show lower difference between saddle and rotary quern grinding than the four-muscle model, as a result of the addition of the muscles presumed to be active during saddle quern grinding (biceps brachii, parts of the deltoid). Finally, based on consistency among our results, we also briefly suggest possible implications of our results for osteological observations, particularly noting a contradiction between our observation of muscle activation and past interpretations of skeletal markers of saddle quern grinding.

## Materials and methods

### Sample

The first sample used in this study was an athletic sample consisting of 10 female rowers (age, 16.3 ± 0.8 years; body height, 170.0 ± 4.8 cm; body mass, 66.4 ± 8.4 kg). According to responses to a questionnaire, our athletic subjects spent on average 10.7 h/week rowing and 2 h/week doing other strenuous tasks involving the upper limb. The second sample consisted of 25 nonathletic females (age, 24.1 ± 2.8 years; body height, 166.3 ± 4.1 cm; body mass, 60.6 ± 9.2 kg) who were recruited mainly from the students of Charles University (Prague, Czech Republic). According to the questionnaire, nonathletic subjects spent on average 3.4 h/week participating in sports and 1.6 h/week doing other strenuous tasks involving the upper limbs. Only right-handed individuals were included in the analyses. Right-handedness was estimated using the method of Oldfield (1971) [22]. Subjects participating in the study gave written consent. The study was approved by the Institutional Review Board of Charles University (Prague, Czech Republic), Faculty of Science (Approval Numbers: 2013/10 and 2017/28).

### Assessment of muscle activation

Muscle activation was assessed using eight surface EMG sensors (Trigno Standard Sensor, Delsys, Natick, MA, USA), which were simultaneously put on the right biceps brachii; anterior, middle, and posterior deltoids; infraspinatus; pectoralis major; and lateral and long heads of the triceps brachii. The skin was cleaned with isopropanol-soaked cosmetic pads [23]. Afterward, sensors were attached to the skin using a double-sided tape (Trigno Sensor Adhesive, Delsys, Inc.). The position and orientation of sensors were according to the recommendation of the Surface Electromyography for the Non-Invasive Assessment of Muscles (SENIAM) project [24]. An exception was the pectoralis major, for which the position of sensor was adopted from Ebben et al. [25], and infraspinatus, for which the position of sensor was adopted from Morris et al. [26]. The EMG signal was acquired with a frequency of 1,926 Hz and filtered using the Butterworth band-pass filter (20–450 Hz) implemented in Delsys Trigno sensors (Delsys, Natick, MA, USA). Raw EMG signal was full wave rectified and smoothed using the root mean square function (window length, 0.125 s; overlap, 0.0625 s) in EMGworks Software (Version 3.21, Delsys, Natick, MA, USA). The mean EMG curve was computed for each subject from at least three subsequent grinding cycles using the Cyclical Analysis script in EMGworks Software. The length of the average cycle was recalculated to 1,000 points using the Resample script in EMGworks Software. The EMG curve values were adjusted to maximum values obtained during maximum voluntary contraction (MVC) tests. To obtain maximum voluntary contractions of all measured muscles, tests were designed for each muscle separately following the recommendations of Konrad [23].

### Grinding

The experimental cereal grinding was performed using the Neolithic saddle quern (Fig 1A) and replica of rotary quern (Fig 1B), which were previously used by Sládek et al. [4]. The saddle quern consists of the lower stationary and mobile upper stone. During saddle quern grinding, the upper stone is pushed back and forth against the lower stone with cereal between them. The rotary quern consists of a stationary lower stone and mobile upper stone, spindle, and handle. During rotary quern grinding, the upper stone is revolved on the lower stone while grain is ground between them.

**Fig 1.**
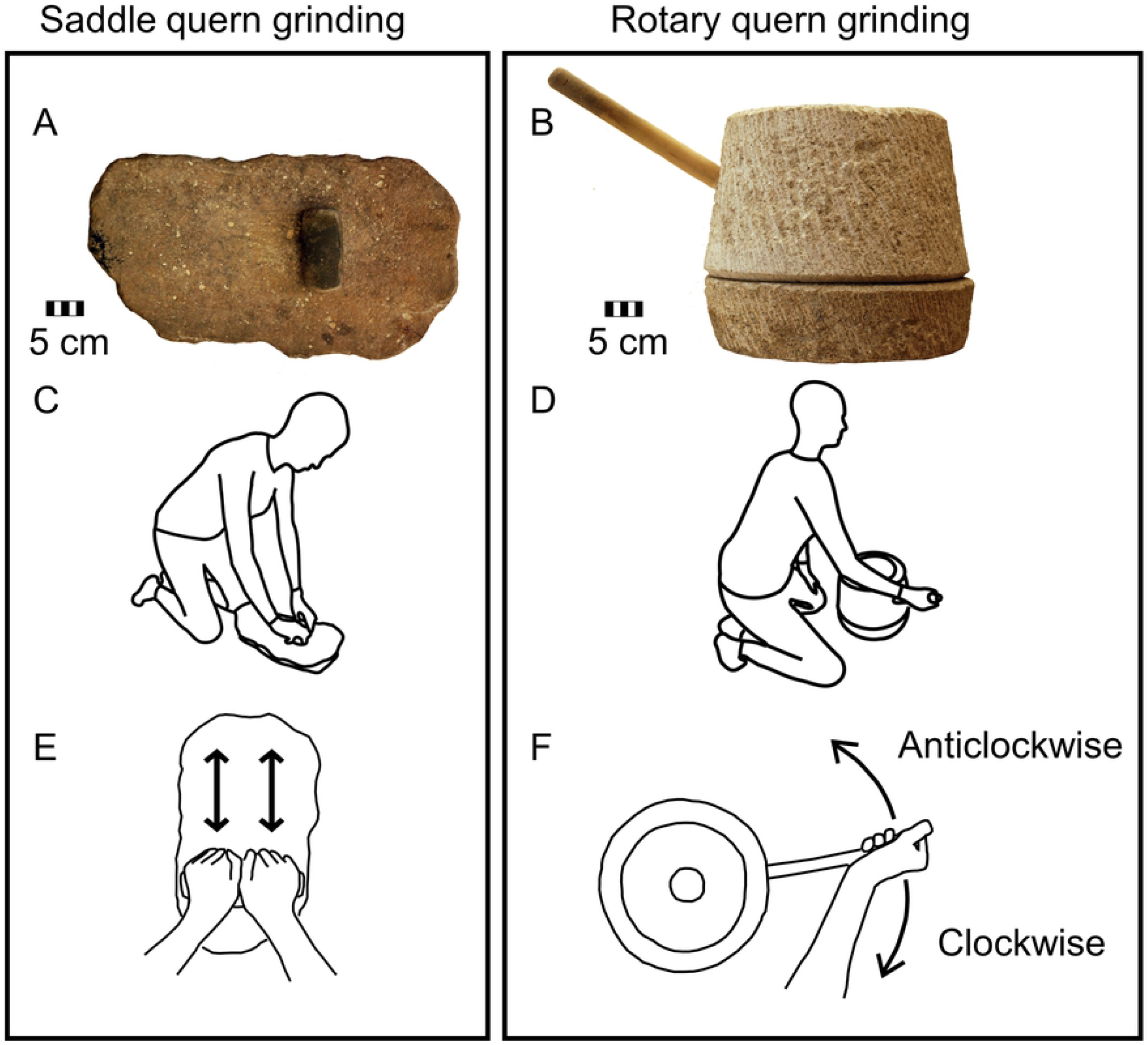
Grinding tools, grinding position, and grinding movements used in the experiment. Saddle quern (A) and rotary quern (B) used in the experiment. Kneeling position used during experimental grinding (C, D). To-and-fro movement used during experimental saddle quern grinding (E) and clockwise and anticlockwise rotary movements used during rotary quern grinding (F).

For both grinding tools, the volunteers were in a kneeling position (Figs 1C and 1D), which was used by Sládek et al. [4]. The kneeling position during saddle quern grinding is supported by lower limb bone alterations in populations, which used saddle querns or grinders [21,27,28], and by ethnographic observation [10]. Rotary querns may be placed on a platform for grinding in a standing position or on the ground for grinding in a kneeling or sitting position [29].

Three experimental grinding tasks were performed, as shown in Figs 1E and 1F. Subjects ground on the saddle quern using both hands in a to-and-fro movement. Saddle quern grinding may also be performed using a circular movement, but the shape of the lower stone used in this study suggests a to-and-fro bimanual movement [4,30]. Rotary quern grinding was performed using the right hand in a circular motion in clockwise and anticlockwise directions. The same movements for saddle and rotary quern grinding were used previously by Sládek et al. [4]. One cycle of saddle quern grinding was defined as one complete to-and-fro motion, and one cycle of rotary quern grinding was defined as complete revolution of the upper stone. Grinding was performed in standardized tempo controlled by metronome, which was 175 and 85 bpm for saddle and rotary quern grinding, respectively. Each grinding cycle was performed at two metronome beats. This grinding tempo was previously used by Sládek et al. [4], where it was selected to be consistent with the preferences of subjects and to maximize grinding effectivity. The different tempo for saddle and rotary quern grinding shouldn’t have an impact on the comparison of athletes and nonathletes, as both samples performed the grinding tasks in the same tempo but could have an impact on the comparison between the saddle and rotary quern grinding. When saddle quern grinding tempo was decreased to 110 bpm, maxEMG increased by 1 % and iEMG increased by 3 %. When clockwise rotary quern grinding tempo was increased to 110 bpm, maxEMG increased by 34 % and iEMG increased by 36 %. Similarly, during anticlockwise rotary quern grinding maxEMG increased by 26 % and iEMG increased by 38 %. If we applied these alterations to previously published results [4], change in tempo would increase the value of the relative difference between the saddle and rotary quern grinding. On the other hand, it wouldn’t influence the pattern of the difference between the saddle and rotary quern grinding, which is a lower muscle activation during saddle than rotary quern grinding [4]. Grinding tasks performed in the present study are shown in S1–S3 Videos.

### Variables

To analyze the magnitude of muscle activation, the EMG curve of each muscle was analyzed using its maximum value (maxEMG, %MVC) and its integral adjusted to the length of cycle (iEMG, %MVC s s^−1^) that described total muscle activity per second. Moreover, the values of iEMG of individual muscles adjusted on physiological cross-sectional area (PCSA) (iEMG_PCSA_, %MVC s s^−1^ cm^2^) were calculated by multiplying iEMG of each muscle by its PCSA. The PCSA values were obtained from a cadaver study [31]. Effectively, iEMG_PCSA_ is an approximate measure of the relative force exerted by the muscle over the course of its activity. Within each subject iEMG_PCSA_ values of all eight muscles were summed to obtain Summed iEMG_PCSA_ of the eight-muscle model, and iEMG_PCSA_ values of the anterior deltoid, infraspinatus, pectoralis major, and long head of the triceps brachii were summed to obtain Summed iEMG_PCSA_ of the four-muscle model. This indicates the combined overall relative force level exerted by all of the included muscles during the measured activity.

The coactivation of muscle pairs was computed using coactivation index as defined by Kellis et al. [32]:

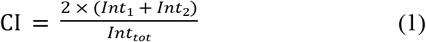

Where Int_1_ and Int_2_ are integrals of EMG signals of two muscles. These integrals were computed in periods, in which the given muscle had lower values of EMG signal than the other muscle. Denominator Int_tot_ is the sum of iEMG of both muscles in a cycle. Coactivation index has values between 0 and 1, with 0 denoting no overlap of EMG signal patterns of the muscles and 1 denoting perfect overlap of EMG signal patterns [32]. The index was computed for each pair of muscles in every subject.

### Statistical analysis

For the purpose of our first goal, we compared maxEMG and iEMG in the athletic and nonathletic samples using the Bonferroni post hoc test as recommended for data violating the assumption of sphericity [33]. Sphericity was analyzed using Mauchly’s sphericity test.

Bonferroni post hoc test was also used to compare coactivation indices between athletes and nonathletes. For our second goal, we compared Summed iEMG_PCSA_ between grinding types within the four- and eight-muscle models in the nonathletic sample using the Tukey HSD test separately within each model. In iEMG_PCSA_ data the assumption of sphericity was met according to Mauchly’s sphericity test. The significance threshold was set to 0.05. Post hoc tests were performed in Statistica for Windows (ver. 12, StatSoft, Inc., 1984–2013).

## Results

The values of maxEMG of athletes and nonathletes are shown in Table 1 and Fig 2. Mean maxEMG was lower in athletes than nonathletes in all muscles, except for the lateral head of the triceps brachii during saddle quern grinding and middle deltoid during clockwise rotary quern grinding. The differences are not significant at the .05 level. The highest difference on average was in posterior deltoid during anticlockwise rotary quern grinding, which had 28.2 %MVC lower maxEMG in athletes than nonathletes (p-value = 0.053). The second highest difference was in the posterior deltoid during clockwise rotary quern grinding, which had 26.2 %MVC (p-value = 0.174) lower mean maxEMG in athletes than nonathletes. On the contrary, the lowest difference between samples occurred in the anterior deltoid during saddle quern grinding, which was 1.1 %MVC less active in athletes than nonathletes. In both samples, the majority of muscles had on average lower maxEMG during saddle quern grinding than rotary quern grinding. An exception was the long head of the triceps brachii, which had higher maxEMG during saddle quern grinding than anticlockwise rotary quern grinding in both samples.

**Fig 2.**
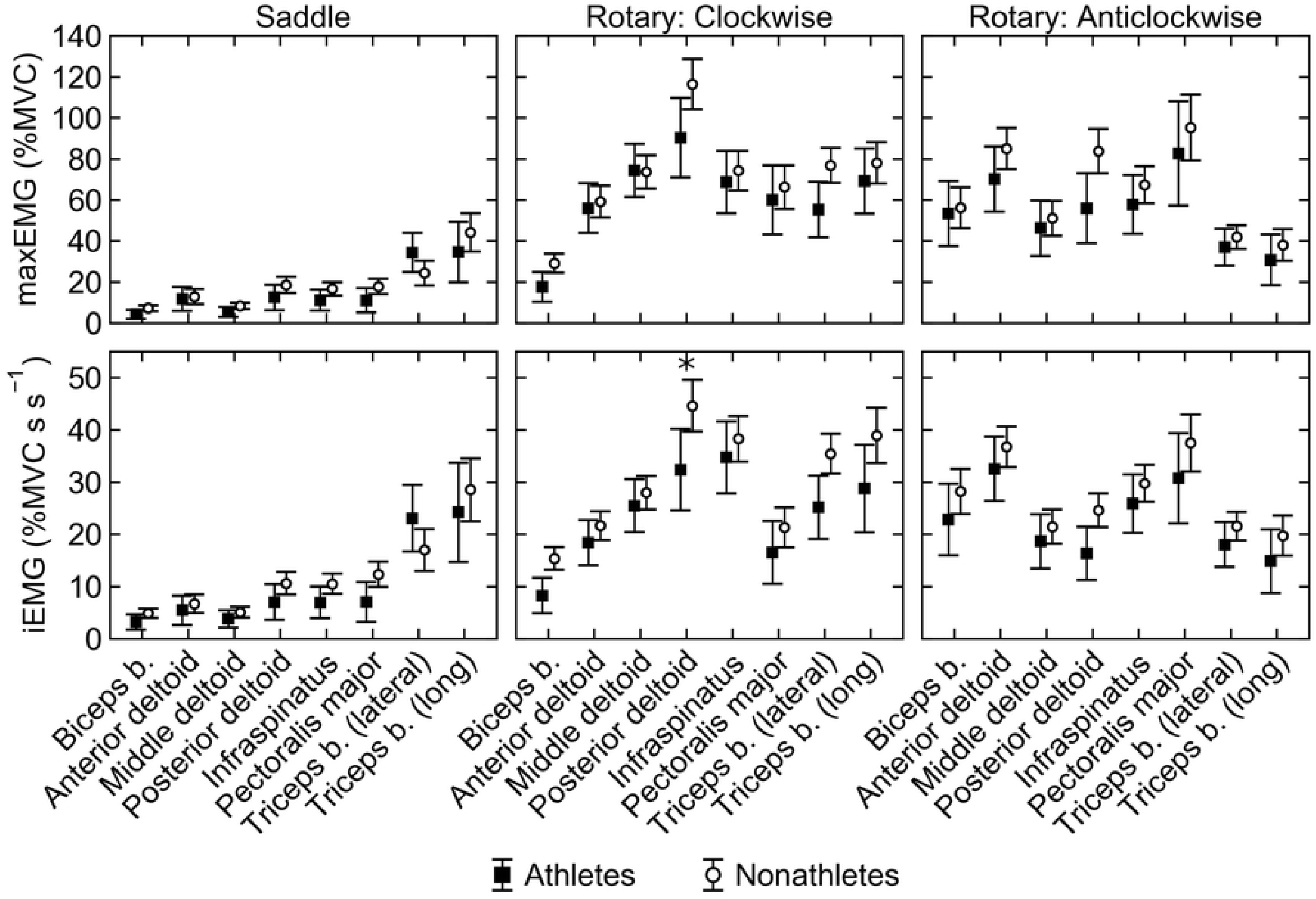
Muscle activation during cereal grinding in athletes and nonathletes. Maximum muscle activation (maxEMG, upper row) and total muscle activity per second (iEMG, lower row) in athletes (black squares) and nonathletes (white circles) during saddle quern (left column), clockwise rotary quern (middle column), and anticlockwise rotary quern (right column) grinding. Markers and whiskers indicate the means and 95% confidence intervals of the means, respectively. Significant difference between athletes and nonathletes (p < 0.05) is indicated by asterisk. See Table 1 for abbreviations of muscles.

**Table 1.** Maximum muscle activation (maxEMG, %MVC).

| **Muscle** | **Saddle** |  | **Rotary: Clockwise** |  | **Rotary: Anticlockwise** |  |
| --- | --- | --- | --- | --- | --- | --- |
|  | **Athletes**<br>**(n = 10)** | **Nonathletes**<br>**(n = 25)** | **Athletes**<br>**(n = 10)** | **Nonathletes**<br>**(n = 25)** | **Athletes**<br>**(n = 10)** | **Nonathletes**<br>**(n = 25)** |
| Biceps b. | 4.2 (2.0–6.5) | 7.2 (5.8–8.6) | 17.5 (10.2–24.8) | 29.1 (24.5–33.7) | 53.4 (37.6–69.1) | 56.4 (46.4–66.3) |
| Anterior deltoid | 11.7 (5.8–17.6) | 12.8 (9.1–16.5) | 55.9 (43.8–68.0) | 59.3 (51.7–66.9) | 70.1 (54.3–85.9) | 85.1 (75.1–95.1) |
| Middle deltoid | 5.4 (3.0–7.9) | 8.3 (6.7–9.8) | 74.3 (61.4–87.2) | 73.7 (65.5–81.8) | 46.2 (32.5–59.8) | 51.4 (42.8–60.1) |
| Posterior deltoid | 12.4 (6.1–18.7) | 18.6 (14.5–22.6) | 90.3 (71.0–109.6) | 116.5 (104.3–128.7) | 55.9 (38.8–73.0)<sup>a</sup> | 84.2 (73.3–95.0)<sup>a</sup> |
| Infraspinatus | 11.0 (5.9–16.2) | 16.7 (13.5–20.0) | 68.7 (53.4–83.9) | 74.3 (64.7–83.9) | 58.1 (43.9–72.3) | 67.6 (58.6–76.6) |
| Pectoralis major | 11.2 (5.2–17.2) | 18.0 (14.2–21.8) | 59.9 (43.1–76.8) | 66.5 (55.9–77.2) | 82.6 (57.2–108.1) | 95.3 (79.2–111.4) |
| Triceps b. (lateral) | 34.8 (25.3–44.3) | 24.5 (18.5–30.5) | 55.3 (41.7–68.8) | 76.9 (68.3–85.5) | 36.9 (27.8–46.0) | 42.0 (36.3–47.8) |
| Triceps b. (long) | 35.1 (20.3–49.9) | 44.3 (35.0–53.7) | 69.2 (53.3–85.1) | 78.0 (68.0–88.1) | 30.7 (18.4–43.0) | 37.9 (30.2–45.7) |

Presented data are mean values with 95% confidence intervals in parentheses. All differences between athletes and nonathletes have p-values > 0.1, except for posterior deltoid during clockwise and anticlockwise rotary quern grinding. P-values are the results of the Bonferroni post hoc test. Biceps b., biceps brachii; triceps b., triceps brachii.
<sup>a</sup> Difference between athletes and nonathletes has p-value = 0.053.

The values of iEMG of athletes and nonathletes are shown in Table 2 and Fig 2. Athletes had lower iEMG than nonathletes in all muscles except for the lateral head of the triceps brachii during saddle quern grinding. The highest mean difference, which was also the only significant one, occurred in the posterior deltoid during clockwise rotary quern grinding, which had 12.3 %MVC s s^−1^ lower mean iEMG in athletes than nonathletes (p-value = 0.035). Differences between samples in other muscles were not significant. The lowest difference between samples was in the anterior deltoid during saddle quern grinding, which was 1.3 %MVC s s^−1^ lower iEMG in athletes than nonathletes. Saddle quern grinding had on average lower iEMG than rotary quern grinding in the majority of muscles in both samples. An exception was the long head of the triceps brachii in both samples and lateral head of the triceps brachii in athletes, which had higher iEMG during saddle quern grinding than anticlockwise rotary quern grinding.

**Table 2.** Total muscle activity per second (iEMG, %MVC s s^−1^).

| **Muscle** | **Saddle** |  | **Rotary: Clockwise** |  | **Rotary: Anticlockwise** |  |
| --- | --- | --- | --- | --- | --- | --- |
|  | **Athletes**<br>**(n = 10)** | **Nonathletes**<br>**(n = 25)** | **Athletes**<br>**(n = 10)** | **Nonathletes**<br>**(n = 25)** | **Athletes**<br>**(n = 10)** | **Nonathletes**<br>**(n = 25)** |
| Biceps b. | 3.2 (1.7–4.6) | 4.9 (4.0–5.8) | 8.3 (4.8–11.7) | 15.4 (13.2–17.5) | 22.8 (15.9–29.7) | 28.2 (23.9–32.5) |
| Anterior deltoid | 5.4 (2.6–8.2) | 6.7 (4.9–8.5) | 18.4 (14.0–22.8) | 21.7 (18.9–24.4) | 32.5 (26.4–38.7) | 36.8 (32.9–40.6) |
| Middle deltoid | 3.8 (2.1–5.4) | 5.1 (4.0–6.1) | 25.5 (20.5–30.5) | 28.0 (24.8–31.1) | 18.6 (13.5–23.8) | 21.5 (18.2–24.7) |
| Posterior deltoid | 7.0 (3.6–10.4) | 10.6 (8.5–12.8) | 32.3 (24.6–40.1)<sup>a</sup> | 44.6 (39.7–49.6)<sup>a</sup> | 16.4 (11.3–21.5) | 24.6 (21.4–27.8) |
| Infraspinatus | 6.9 (3.9–10.0) | 10.5 (8.6–12.4) | 34.8 (27.9–41.7) | 38.3 (33.9–42.7) | 25.9 (20.3–31.5) | 29.8 (26.2–33.3) |
| Pectoralis major | 7.0 (3.2–10.8) | 12.3 (9.9–14.7) | 16.5 (10.5–22.6) | 21.3 (17.5–25.1) | 30.7 (22.1–39.4) | 37.5 (32.0–43.0) |
| Triceps b. (lateral) | 23.1 (16.7–29.4) | 17.0 (13.0–21.0) | 25.2 (19.1–31.2) | 35.4 (31.6–39.3) | 18.0 (13.7–22.3) | 21.6 (18.8–24.3) |
| Triceps b. (long) | 24.2 (14.7–33.7) | 28.5 (22.5–34.5) | 28.8 (20.4–37.1) | 38.9 (33.6–44.2) | 14.8 (8.7–21.0) | 19.7 (15.9–23.6) |

<sup>a</sup> Difference between athletes and nonathletes has p-value = 0.035.

The differences of coactivation index between athletes and nonathletes are shown in S1–S3 Tables. Higher antagonistic coactivation could indicate less efficient movement strategy, during which the agonistic muscle must produce more force to overcome moment of force of the antagonistic muscle. Coactivation indices were lower in athletes than nonathletes in 64 out of 84 comparisons. Most of the differences between athletes and nonathletes were not significant. The only significant difference between samples occurred in the coactivation index between the lateral head of the triceps brachii and pectoralis major during saddle quern grinding, when athletes had on average 30 % lower (p-value = 0.048) coactivation index than nonathletes.

Summed iEMG_PCSA_ of the eight- and four-muscle models is shown in Table 3. By comparing Summed iEMG_PCSA_ of saddle and rotary quern grinding using the eight- and four-muscle models, we can show whether the larger set of muscles changes the estimates of the upper limb loading during saddle and rotary quern grinding. Saddle quern grinding had on average significantly lower Summed iEMG_PCSA_ than rotary quern grinding (in either direction) in both models. Summed iEMG_PCSA_ during saddle quern grinding was on average 2.7 times lower than that during clockwise rotary quern grinding (p-value = 0.0001) in the eight-muscle model and 2.3 times lower (p-value = 0.0001) in the four-muscle model. Summed iEMG_PCSA_ during saddle quern grinding was on average 2.6 times lower than that during anticlockwise rotary quern grinding (p-value = 0.0001) in the eight-muscle model and 2.5 times lower (p-value = 0.0001) in the four-muscle model. In the four-muscle model, Summed iEMG_PCSA_ during clockwise rotary quern grinding was non-significantly lower than that during anticlockwise rotary quern grinding (p-value = 0.066). On the contrary, in the eight-muscle model, Summed iEMG_PCSA_ during clockwise rotary quern grinding was non-significantly higher than that during anticlockwise rotary quern grinding (p-value = 0.354).

**Table 3.** Total activity of all measured muscles adjusted to PCSA (Summed iEMG_PCSA_, %MVC s s^−1^ cm^2^) in the eight- and four-muscle models.

| Type of grinding | Eight-muscle model (n = 25) | Four-muscle model (n = 25) |
| --- | --- | --- |
| Saddle | 507.9 (435.2–580.6)<sup>a,b</sup> | 334.5 (279.5–389.5)<sup>a,b</sup> |
| Rotary: Clockwise | 1376.7 (1282.1–1471.4)<sup>c</sup> | 763.7 (697.5–829.9)<sup>c</sup> |
| Rotary: Anticlockwise | 1303.8 (1179.2–1428.4)<sup>c</sup> | 847.5 (763.6–931.4)<sup>c</sup> |

Presented data are mean values with 95% confidence intervals in parentheses. Significance was determined using the Tukey HSD test.
<sup>a</sup> Significantly different from clockwise rotary quern grinding.
<sup>b</sup> Significantly different from anticlockwise rotary quern grinding.
<sup>c</sup> Significantly different from saddle quern grinding.

## Discussion

Our first goal was to compare muscle activation during cereal grinding between athletes and nonathletes. We expected athletes to have lower muscle activation than nonathletes during grinding due to lower antagonistic coactivation in athletes. While there was a trend of lower activation in athletes than in nonathletes, most differences in activation were not significant. Similarly, coactivation was lower in athletes than nonathletes in most muscle pairs, but significant difference occurred only in one muscle pair, which was not antagonistic. Our results therefore suggest that nonathletic subjects can be used for reconstruction of saddle and rotary quern grinding. Furthermore, the broader test of the relative muscle activity in saddle versus rotary quern grinding is fully consistent; in both samples rotary quern grinding involves significantly more muscle activation than saddle quern grinding. Our second goal was to test the influence of the choice of analyzed muscles on the estimate of upper limb loading during cereal grinding. Again, both the eight- and four-muscle models indicated that the upper limb is loaded significantly less during saddle quern grinding than rotary quern grinding, which suggests that the upper limb muscles may be reduced to the four-muscle model for comparison of saddle and rotary quern grinding.

### Muscle activation during grinding in athletes and nonathletes

There was a trend of lower average maxEMG and iEMG in athletes than nonathletes, in which athletes had on average 16 % lower maxEMG and 19 % lower iEMG than nonathletes during cereal grinding. On the other hand, significant differences between athletes and nonathletes occurred only in the posterior deltoid during clockwise rotary quern grinding. Furthermore, both athletes and nonathletes had lower activation during saddle quern grinding than rotary quern grinding in the majority of muscles, supporting the previous analysis based on nonathletes.

Lower activation of the upper limb muscles in our athletic than nonathletic sample could be caused by lower antagonistic coactivation, higher muscle activation in other parts of the body, or higher muscle hypertrophy in athletes than nonathletes. Antagonistic coactivation would have an impact on both muscle activation and force because with lower coactivation, the agonistic muscle would have to produce less force to overcome the force of the antagonistic muscle. Even though in most muscle pairs coactivation was on average lower in our athletic than nonathletic sample, no significant difference was noted between samples in the coactivation of the antagonistic muscles. The lack of significant differences in coactivation between athletes and nonathletes could be also associated with similar timing of muscle activation in both samples (Fig 3). In any case, the similar levels of antagonistic coactivation in athletes and nonathletes support the use of nonathletes in experimental cereal grinding.

**Fig 3.**
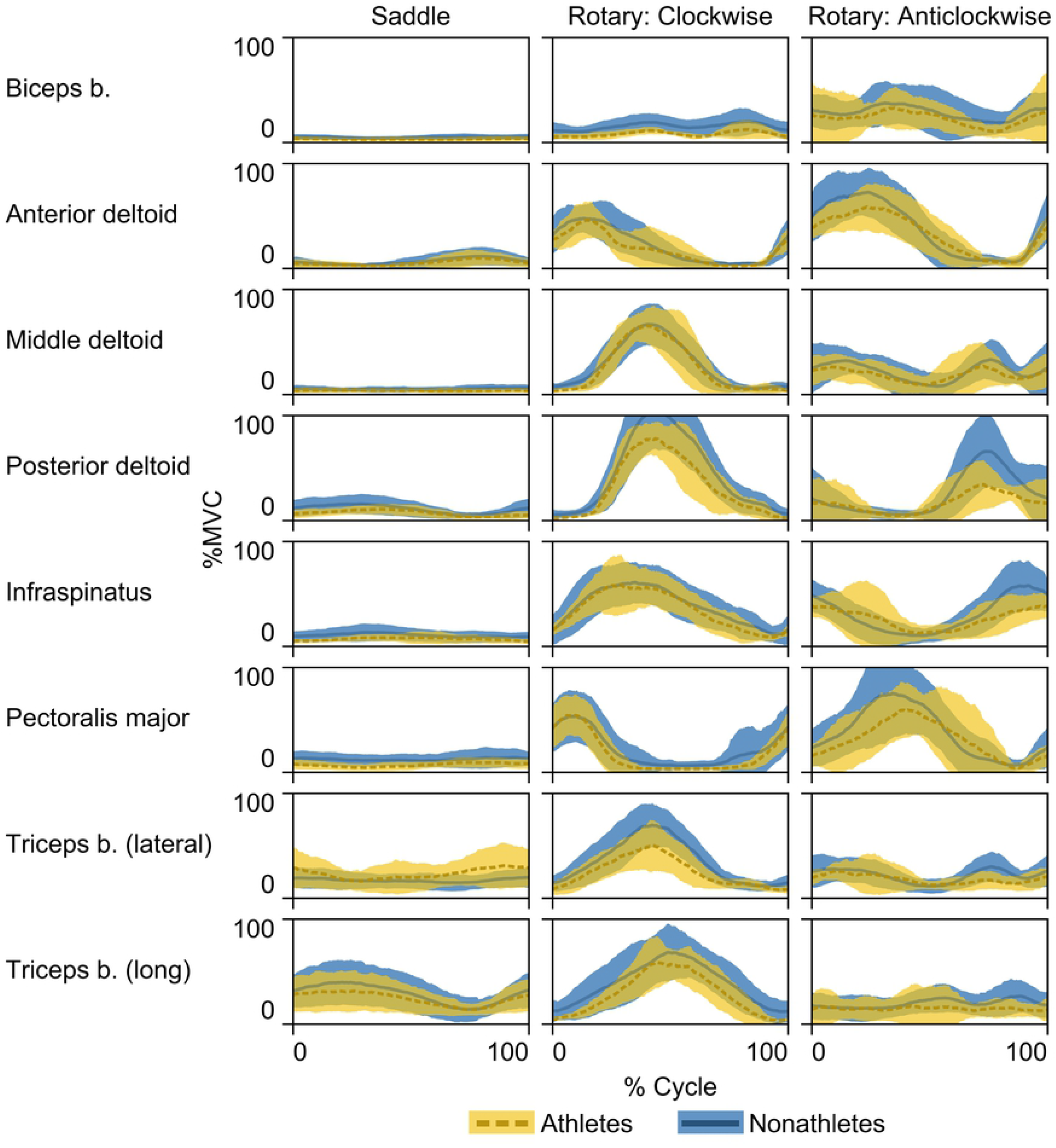
Averaged EMG curves. Results are shown for athletic (yellow, dashed line) and nonathletic (blue, solid line) samples during saddle quern (left column), clockwise rotary quern (middle column), and anticlockwise rotary quern (right column) grinding. Darker lines and colored areas indicate the means and ± standard deviations, respectively. See Table 1 for abbreviations of muscles.

Another possibility is that lower activation of the upper limb muscles in our athletic than nonathletic sample was caused by the activation of the muscles of other parts of the body than the upper limbs. For example, it was observed that Hopi women used the whole body to produce rhythmic movements during grinding on saddle quern [34]. Rowers may be able to activate the muscles of the whole body better than nonathletes because during rowing they generate power with the upper and lower limbs and torso [35]. Therefore, we cannot rule out the possibility that lower activation of the upper limb muscles in athletes was caused by their higher activation of the muscles in other parts of the body. However, again it is notable that differences between athletes and nonathletes are largely insignificant, with the exception of a single muscle (posterior deltoid), and the direction of contrast between saddle and quern grinding was consistent between the two samples.

Alternatively, lower activation of the upper limb muscles in athletes than nonathletes may be caused by increased muscle hypertrophy in athletes, indicating different muscle force between athletes and nonathletes [36,37]. It was observed that with increasing muscle hypertrophy, activation per muscle force decreases [37]. Therefore, a hypertrophied muscle may produce the same muscle force with lower muscle activation than a muscle without hypertrophy. Therefore, if increased hypertrophy was the reason for lower muscle activation in athletes than nonathletes, this difference would not have an impact on muscle force stressing the upper limb bones. Muscle hypertrophy can be increased by both resistance training [38] and endurance training [39]. Therefore, it may be expected in the upper limb muscles of rowers. Rowing power output was shown to be correlated with hypertrophy of the muscles that extend the elbow [35], which suggests that those muscles could be hypertrophied in our athletic sample as well. Therefore, we cannot rule out the possibility that lower muscle activation in athletes was offset by hypertrophied muscles in athletes. As hypertrophy only influences muscle activation and not muscle force, athletes would produce the same muscle force as nonathletes. Therefore, either of the samples may be used for the estimation of upper limb loading during grinding, if muscle hypertrophy is considered.

Our results might suggest that influence of athletic samples is limited to reconstructions of tasks in which the athletes are specialized, such as reconstruction of spear throwing using javelin throwers [6]. Furthermore, the difference between athletes and nonathletes might present itself only in some variables describing performance, such as success rate during spear throwing [6], while other measures, such as muscle activation, might be similar in athletes and nonathletes. Future studies of cereal grinding might therefore compare athletes with nonathletes in other measures, such as grinding efficiency.

### Influence of analyzed muscles on the estimated upper limb loading

Upper limb muscle loading was significantly lower during saddle quern grinding than rotary quern grinding in both eight- and four-muscle models. The similarity of results from both models for comparison of saddle and rotary quern grinding supports reducing the upper limb muscles to the four-muscle model.

In our results, both eight- and four-muscle models indicate that the upper limb is less loaded during saddle quern grinding than rotary quern grinding. However, the eight- and four-muscle models differed in comparison of upper limb loading during clockwise rotary quern grinding and the other two grinding types. The eight-muscle model suggested higher difference in loading between clockwise rotary and saddle quern grinding than the four-muscle model. Observed difference between the eight- and four-muscle models may be explained by the muscles included in models and their importance for either grinding type. During saddle quern grinding, the highest iEMG_PCSA_ was in the pectoralis major and long head of the triceps brachii, which were included in both models. During anticlockwise rotary quern grinding, the highest iEMG_PCSA_ was in the anterior deltoid, infraspinatus, and pectoralis major, which were also included in both models. The inclusion of these muscles in both models perhaps led to similar ratio of Summed iEMG_PCSA_ of anticlockwise rotary and saddle quern grinding in both models. During clockwise rotary quern grinding, the second and third highest iEMG_PCSA_ were in the posterior and middle deltoid, respectively. As these two muscles were only in the eight-muscle model, their inclusion may have increased the ratio between Summed iEMG_PCSA_ of clockwise rotary and saddle quern grinding in the eight-muscle model. Therefore, the four-muscle model may underestimate upper limb loading during clockwise rotary quern grinding relative to saddle quern grinding. Nevertheless, both models showed significantly lower Summed iEMG_PCSA_ in saddle quern grinding than rotary quern grinding. Therefore, the upper limb muscles could be reduced to four muscles for comparison of upper limb loading during saddle and rotary quern grinding.

In our results, the most active muscle during saddle quern grinding was the long head of the triceps brachii, which had the highest values of activation (maxEMG and iEMG) and the second highest activation adjusted to PCSA (iEMG_PCSA_; Fig 4). The long head of the triceps brachii was also the only muscle that had higher maxEMG and iEMG during saddle quern grinding than rotary quern grinding. High values of maxEMG and iEMG of the long head of the triceps brachii were also observed in a previous cereal grinding study [4]. High activation of the triceps brachii relative to other muscles during saddle quern grinding shows the possibility of using its origin and insertion in future studies of entheseal changes associated with saddle quern grinding. Saddle quern grinding has previously been suggested to cause entheseal changes of deltoid insertion [21]. In our results, the values of iEMG_PCSA_ of each deltoid part were on average lower than those of the most active muscles, but the combined value of all deltoid parts would be the second highest of the analyzed muscles. Saddle quern grinding was also previously suggested to be the cause of prominent insertion of the biceps brachii [21]. Therefore, it would be expected to have high values of activation. On the contrary, our results showed that the biceps brachii had the lowest maxEMG, iEMG, and iEMG_PCSA_ of measured muscles during saddle quern grinding. During rotary quern grinding, the infraspinatus and pectoralis major had the highest iEMG_PCSA_; thus, their insertions could be candidates for entheseal changes associated with rotary quern grinding. Nevertheless, if iEMG_PCSA_ values of the anterior, middle, and posterior deltoids are combined, the resulting value is higher than other muscles’ iEMG_PCSA_ during rotary quern grinding; thus, the deltoid may be significant for rotary quern grinding as well. For further research of physical activity during adoption of agriculture, the pectoralis major, triceps brachii, and previously suggested deltoid may be candidate muscles for analysis of entheseal changes. For studies of physical activity during intensification of agriculture, the deltoid muscles, infraspinatus, and pectoralis major may be candidate muscles for analysis of entheseal changes.

**Fig 4.**
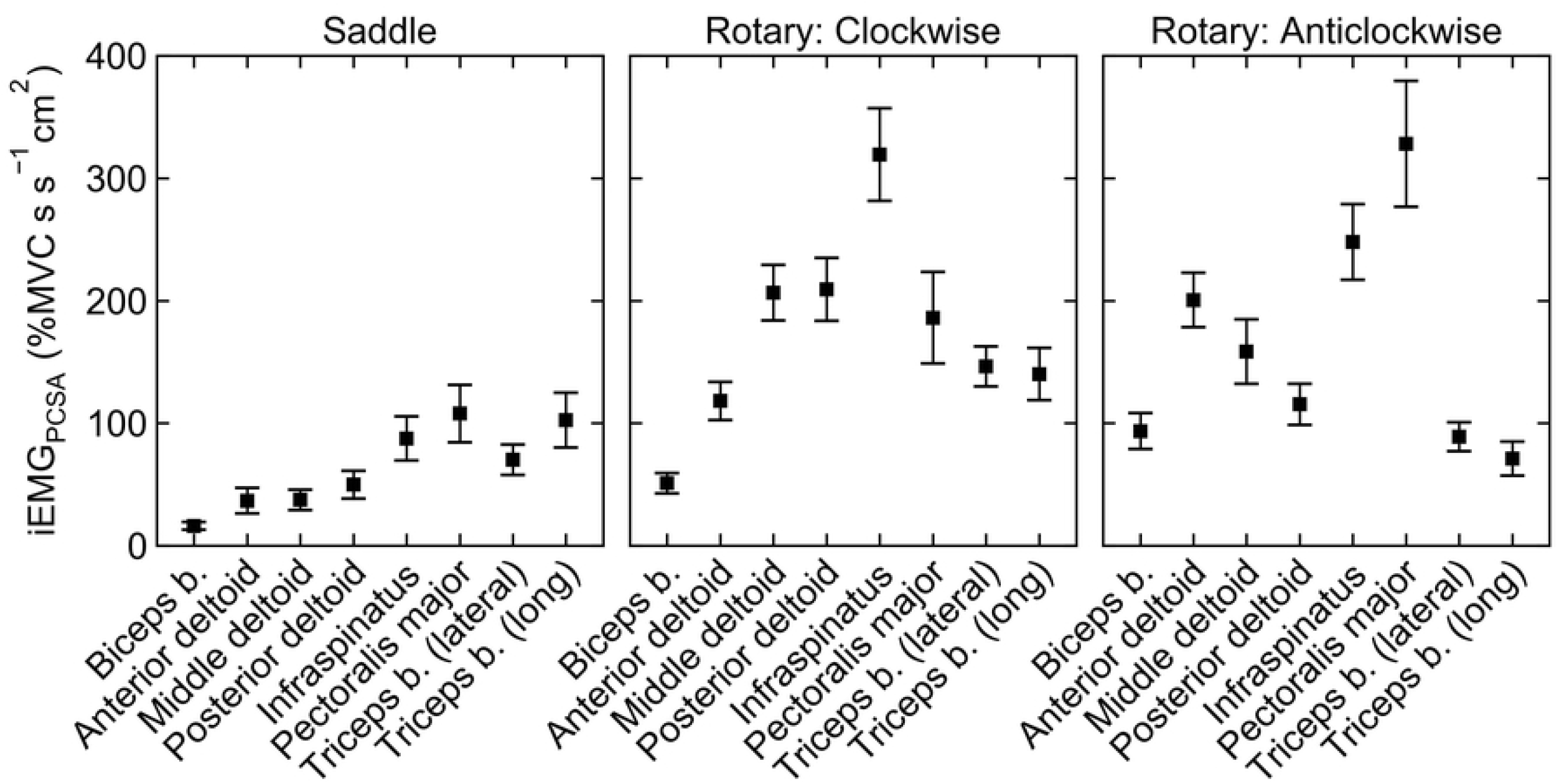
Total muscle activity per second adjusted for PCSA of the muscles (iEMG_PCSA_) in nonathletic sample. Results are shown for saddle quern (left), clockwise rotary quern (middle), and anticlockwise rotary quern (right) grinding. Markers and whiskers indicate the means and 95% confidence intervals of the means, respectively. See Table 1 for abbreviations of muscles.

## Conclusions

Our results showed a trend of lower activation in the muscles of athletes relative to nonathletes; however, these differences mostly were not significant. Furthermore, both athletes and nonathletes had lower activation during saddle quern grinding than rotary quern grinding in the majority of muscles. Our results therefore indicate that both athletes and nonathletes can be used for comparison of upper limb loading during saddle and rotary quern grinding. Our results also suggest that a homogenous sample would be favorable for estimation of muscle force during cereal grinding to minimize influence of subjects’ muscle hypertrophy on muscle activation.

Both eight- and four-muscle models showed lower upper limb loading during saddle quern grinding than rotary quern grinding, which suggests that the upper limb muscles may be reduced to the previously used four-muscle model for comparison of saddle and rotary quern grinding.

Finally, our results also suggest refinements in the use of osteological markers of entheseal changes to analyze physical activity changes during the adoption of agriculture. These changes associated with grinding should be observed for the deltoid, infraspinatus, and pectoralis major for both saddle and rotary quern grinding or for the long head of the triceps brachii for saddle quern grinding. However, the low level of biceps brachii involvement raises questions about the interpretation of prominent radial tuberosity development as specifically related to saddle quern grinding [21] and suggests that alternative strenuous activities might be considered as responsible for this skeletal indicator.

More generally, our results confirm that experimental replicative research on activity patterns involving modern populations may yield fairly robust results, despite the contrast in level of conditioning of most modern populations, and with a judiciously chosen, limited set of monitored muscles. Furthermore, it offers the possibility of more broadly examining muscle activation in a range of prehistoric activities to better assess and test interpretations of prehistoric skeletal modifications.

## Acknowledgments

We would like to thank the participants from Charles University and from the rowing teams Český veslařský klub Praha, Veslařský klub Smíchov, and SK HAMR. We also wish to thank the coaches and staff of the participating rowing teams. We would like to thank Iva Brynychová, Pavla Alexia Jarešová, Klára Kosatíková, and Zuzana Matějovská for their help with data collection. We would also like to thank Jakub Otáhal for the advice about EMG data collection.

## Supporting information captions

**S1 Video. Cereal grinding using saddle quern.** Synchronized acquisition of electromyography data is shown in the right part of the video. Electromyography data was full wave rectified and smoothed using root mean square function. See Table 1 for abbreviations of muscles.

**S2 Video. Cereal grinding on rotary quern in clockwise direction.** Synchronized acquisition of electromyography data is shown in the right part of the video.

Electromyography data was full wave rectified and smoothed using root mean square function. See Table 1 for abbreviations of muscles.

**S3 Video. Cereal grinding on rotary quern in anticlockwise direction.** Synchronized acquisition of electromyography data is shown in the right part of the video.

Electromyography data was full wave rectified and smoothed using root mean square function. See Table 1 for abbreviations of muscles.

**S1 Table. Coactivation indices during saddle quern rotary quern grinding in athletes and nonathletes.**

**S2 Table. Coactivation indices during clockwise rotary quern grinding in athletes and nonathletes.**

**S3 Table. Coactivation indices during anticlockwise rotary quern grinding in athletes and nonathletes.**

